# Differences between protein fitness models can be used to design variants of altered specificity

**DOI:** 10.64898/2026.06.10.731299

**Authors:** Samuel P. Berry, Rachelle Gaudet, Debora S. Marks

## Abstract

The vastness of sequence space makes it challenging for directed evolution to efficiently traverse the fitness landscape. In recent years, unsupervised probabilistic models trained on natural sequences have shown promise for predicting the functional effects of mutations and designing new proteins, leading many to suggest that these models may be useful for guiding directed evolution campaigns toward functional regions of sequence space. However, many directed evolution campaigns are interested in evolving new substrate or ligand specificity, and the behavior of unsupervised sequence models on predicting and designing activity against altered substrates or ligands has not been tested. We have built a curated database of multiplexed functional assay results profiling substrate or ligand specificity and systematically how assessed various popular unsupervised protein fitness models score and design variants that alter selectivity in these datasets. We find that sequence models that learn from the surrounding sequence context, especially protein language models, systematically bias against variants that alter specificity. This bias leads them to design altered-specificity variants at similar or lower rates than random chance. However, we propose a simple strategy to exploit this bias by taking weighted differences of model scores to enrich libraries for altered-specificity variants. These findings and our database should help guide how biologists best use protein fitness models and provide a framework to help machine learning researchers develop a new generation of machine learning models that can better design novelty.

## Introduction

The rapidly developing ability to engineer new proteins via directed evolution has revolutionized biology with massive biotechnological implications in health, industry, energy, climate change, and other fields^1, 2^. A common task is to take a natural protein with a defined function and evolve an alteration to that function, for example a change to the substrate or ligand the protein recognizes. However, directed evolution is often limited by the vast scope of sequence space and the impossibility of testing even a fraction of the total possible sequences^3, 4^. Thus, many new protein functions that could be theoretically encoded in a short amino acid sequence remain undiscovered. While in theory all a protein’s functional properties could be predicted from its amino acid sequence using chemical and physical principles, in practice this approach is computationally intractable. By contrast, scientists have had far more success predicting the functional properties of proteins represented in large natural sequence databases, leveraging the assumption that observed sequences continue to exist because they are fit. This evolutionary framework underlies successful methods for predicting protein structure^5, 6^, identifying stabilizing mutations that preserve function^7, 8^, and predicting the clinical effects of mutations^9, 10^.

These models have been systematically evaluated by comparing their predictions against protein “fitness” scores obtained from deep mutational scans^11^. Strikingly, initial results suggested that even relatively simple models trained on multiple sequence alignments (MSAs) such as a Potts model^6, 12^ could predict these experimental results with reasonable accuracy, despite enormous differences in how these experiments read out protein function. Since then, many new machine learning models have the explicit goal of predicting protein fitness, with notable new classes including protein language models and inverse folding models (see Supplemental Note 1 for a description of model classes). We continually update a database of models which are assessed against hundreds of deep mutational scans as ProteinGym^13^, which has a leaderboard of models with highest average correlation across all experiments in the database.

The ability of these models to predict scores from deep mutational scans is often related to an ability to design new protein variants. Indeed, several probabilistic models trained purely on natural sequence data are able to generate sequences with similar functions to a native sequence^14–18^. Some designed variants have improved thermostability or other desirable properties^19, 20^, suggesting that these models may be useful for protein engineering. Some studies have concluded that these models learn a generalized notion of protein fitness and therefore that protein fitness models may be useful for guiding evolution campaigns, even for functions distinct from those of the native protein^17, 21–24^. Protein language models have emerged as the most popular tool for this task^25^. However, there has been no systematic evaluation of the usefulness of either mutational effect prediction or protein design with generative probabilistic models for redesigning natural proteins for altered specificity, despite the broad interest of this question in the directed evolution field.

There is additional ambiguity about how protein fitness models behave when trained on many sequences of altered function. Unsupervised probabilistic models fit a single probability distribution over the entire data distribution, even though natural protein sequences evolved under diverse selective pressures. For instance, even within an alignment of a single family of protein sequences the fitness landscapes represented can be very heterogeneous, from subtle changes in organismal niche to major changes in the evolutionarily selected specificity or regulation of paralogs. The challenge of training set heterogeneity is especially true for protein language models, which are trained on all known protein sequences—and thus functions—at once. We propose two potential hypotheses for how a single model trained on such heterogeneous landscapes would behave. The first is that models trained on all natural sequences could have learned general rules about what makes a functional protein^22^, and thus could design proteins agnostically to their particular function as long as they are functional. If so, these models should assign similar scores to mutations that preserve a protein’s native function and ones that introduce a new function. The second hypothesis is that these models fit very closely to the natural distribution and learn an implicit mixture of family- and function-specific “modes”^26^ rather than learning generalizable rules. In this case, deep learning models would only assign high log-likelihoods to mutations that preserve a native protein’s function, selecting against variants that alter that native function even if these variants are functional along a new axis.

Here we build a companion database to ProteinGym called SpecificityStudio, in which we focus on how different generative models score mutations that alter specificity among a small, curated set of deep mutational scans. We find that none of the major current models are broadly useful for enriching directed evolution libraries for mutations that alter specificity, but that different classes of models treat specificity-altering mutations very differently. Finally, we leverage these differences to build ensemble models that *do* enrich for specificity-altering mutations. Our findings can guide a new generation of probabilistic sequence models that are specificity-aware and capable of biasing toward novelty.

## Results

### A benchmark set of mutational scans for altered specificity

We collected a diverse benchmark set of eight multiplexed functional assay datasets that each profile a large set of variants against multiple substrates or ligands with distinct properties (Table 1 and Figure 1). These datasets vary considerably in all aspects. The proteins assayed include a binding domain (PSD95-PDZ3), three enzymes (TEM-1, AmiE, PafA), two transcription factors (Pho4, RamR), and two transporters (NorA, DraNramp). In six cases, the variants profiled correspond to all single mutations to the wildtype protein, while others have combinatorial variants at a set of active-site residues or a combination of single and combinatorial mutants. The nature of each functional assay also differs. In the PafA and Pho4 studies, variants were purified on a microfluidic device to enable the measurement of precise biochemical constants—the catalytic efficiency (kcat/K_M_) for PafA^27^ and the free energy of binding for Pho4^28^. However, these studies are more limited in their library size. All other studies are done in cells, with some reaching very large scales (over 10^5^ measurements for PSD95-PDZ3^29^ and RamR^30^) but with less precision about particular biochemical parameters. For some datasets, such as the multidrug efflux pump NorA^31^, all profiled substrates are native to the wildtype protein. For others, such as the selective metal importer DraNramp^32^, the alternate substrate is completely excluded by the wildtype protein.

**Table 1.** The eight datasets in SpecificityStudio v1.0.

| Name | Protein group | Native function | Assayed substrates/ ligands | Dataset class* | No. target variants | Mutational depth | Measurement |
| --- | --- | --- | --- | --- | --- | --- | --- |
| **PSD95-PDZ3**<sup>29, 59</sup> | Binding domain | Binds C-terminal peptides such as CRIPT | Single mutants to CRIPT ligand | 1 | 1,772 | Only single mutants | BindingPCA /AbundancePCA |
| **TEM-1**<sup>34</sup> | Enzyme (serine hydrolase) | Hydrolyzes β-lactam antibiotics | Ampicillin and cefotaxime | 1 | 4,997 | Only single mutants | Bacterial growth |
| **AmiE**<sup>60</sup> | Enzyme (amidase) | Hydrolyzes short aliphatic amides | Three separate amides of varying chain length | 2 | 6,820 | Only single mutants | Bacterial growth |
| **PafA**<sup>27</sup> | Enzyme (phosphatase) | Hydrolyzes phosphate monoester | Two phosphate monoesters (MeP and cMUP) | 2 | 1,042 | Only single mutants | $k_{cat}/K_M$ (HT-MEK) |
| **Pho4**<sup>28</sup> | Transcription factor | Binds DNA sequence CACGAG to activate transcription | Single variants to cognate DNA sequence | 1 | 215 | Only single mutants | $\Delta G_{binding}$ (STAMMPing) |
| **RamR**<sup>30</sup> | Allosteric transcription factor | Depresses transcription upon binding ligands | Two enantiomeric drug precursors (1S-THIQ and 1R-THIQ) and their achiral precursor (1-DHIQ) | 1 | 139,438 | 1-12 mutants, mostly 2-6 | Transcriptional induction tied to bacterial growth |
| **NorA**<sup>31</sup> | Multidrug exporter | Promiscuously exports many small molecules | Eight diverse small molecules, all natively exported by NorA | 2 | 7,344 | Only single mutants | Bacterial growth |
| **DraNramp** | Transporter | Selectively imports $Mn^{2+}$ and $Fe^{2+}$ | $Mn^{2+}$ and $Mg^{2+}$ | 1 | 10,229 | 1-10 mutants, mostly 1-4 | Bacterial reporter assays |

\* Class I datasets profile specificity shifts toward new substrates or ligands, and Class II datasets profile specificity shifts among existing substrates or ligands (Supplemental Note 2).

**Figure 1.**
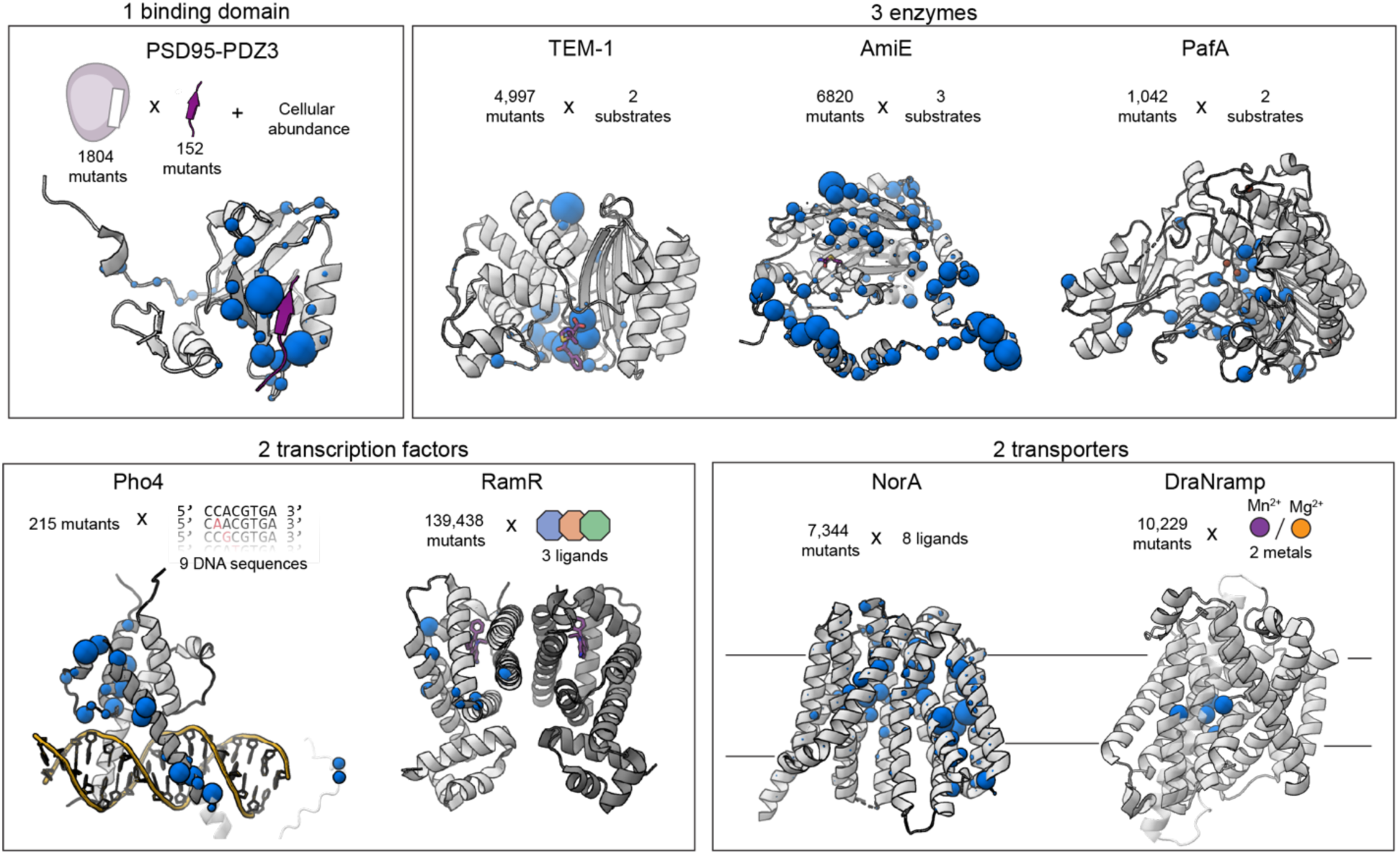
SpecificityStudio: a curated collection of mutational scans for altered specificity. Structures of all proteins with datasets included in the initial version of SpecificityStudio, grouped by protein type. Positions with specificity-altering mutations are highlighted as blue spheres and sized according to the number of specificity-altering single mutations at that position. The PDB codes used are 5HEB^69^ (PSD95-PDZ3), 1M40^70^ (TEM-1), 3LAT^71^ (AmiE), 5TJ3^72^ (PafA), 1A0A^73^ (Pho4), 3VVY^74^ (RamR), 9B3O^75^ (NorA), and 8E60^76^ (DraNramp). Pho4 has several specificity-altering mutations in disordered regions beyond what is resolved in the crystal structure; to indicate this, a partially transparent AlphaFold2 structure is overlaid.

The variable nature of the studies necessitates variable strategies to identify specificity-altering mutations. This process is simplest for the two based on *in vitro* assays, as in these cases the effects for different substrates are nearly linearly related with some outliers that can be defined by a simple linear regression approach (Supplemental Figure 1). Due to the nonlinear mapping between biochemical energies and observed cell-based parameters, this approach does not always work as well for the cell-based assays. Zarin and Lehner^29^ use a similar regression approach but account for global epistasis by fitting a MoCHI energy model^33^ with sigmoidal global epistasis. They assume additive effects between substrate and protein mutations, then identify statistical deviations from additivity that represent specificity. For TEM-1^34^, RamR^30^ (Supplemental Figure 2), and DraNramp^32^, the wildtype protein had little or no activity on the altered-specificity ligands or substrates in question, so we simply identify altered-specificity variants as those that had any activity on these alternate substrates.

### Context-aware fitness models bias against altered-specificity variants

For each dataset, we separated mutations into one of three classes: one representing a strong signal of complete inactivity (“inactive”), one representing functional proteins with similar specificity to the wildtype protein (“native specificity”) and one representing generally functional proteins with an altered specificity profile (“altered specificity”). The exact definition of these classes varies by dataset due to the distinct nature of the data and is presented in Supplemental Note 2 and Supplemental Figures 1-2. We then scored all variants in each dataset with one of fourteen probabilistic fitness models from one of six classes (Table 2).

**Table 2.** The 14 protein fitness models in SpecificityStudio v1.0.

| Name | Class | Trained on | Architecture | Notes |
| --- | --- | --- | --- | --- |
| **Site-independent (PSSM)** | Conservation | MSA | PSSM | From EVmutation pipeline |
| **EVmutation<sup>11</sup>** | Alignment covariation | MSA | Potts model | Inferred with pseudolikelihood maximization |
| **GEMME<sup>56</sup>** | Alignment covariation | MSA | Phylogeny-based | Run with Docker container |
| **EVE<sup>9</sup>** | Alignment covariation | MSA | Variational autoencoder | Ensemble of five models |
| **ESM-1v<sup>37</sup>** | Single-sequence (PLM) | Uniref90 (2020.03) | Masked language model | Ensemble of five models |
| **ESM-2<sup>61</sup>** | Single-sequence (PLM) | Uniref50 + Uniref90 (2022.08) | Masked language model | Single model |
| **ProGen2<sup>62</sup>** | Single-sequence (PLM) | Uniref90 + BFD30 | Autoregressive language model | Used XL model |
| **Tranception (no retrieval)<sup>63</sup>** | Single-sequence (PLM) | UniRef100 | Autoregressive language model | Used L model |
| **MSA Transformer<sup>64</sup>** | Hybrid alignment-based + PLM | Uniref50 (2018.03) and MSA | Masked language model |  |
| **PoET<sup>65</sup>** | Hybrid alignment-based + PLM | Uniref50 clusters and MSA | Autoregressive language model |  |
| **ProteinMPNN<sup>66</sup>** | Inverse folding | PDB (2021.08) | Graph neural network | Predictions made on an AlphaFold prediction of the WT sequence |
| **ESM-IF1<sup>67</sup>** | Inverse folding | CATH and Uniref50 (2022.08) | Masked language model with graph neural network | Predictions made on an AlphaFold prediction of the WT sequence |
| **SaProt<sup>68</sup>** | Hybrid: PLM with structural tokens | Uniref90 (2022.08) + 40M AlphaFold2 models | Masked language model | 650M parameter model trained on AlphaFold2 |
| **ESCOTT<sup>57</sup>** | Hybrid: MSA with structural tokens | MSA, input structure | Phylogeny-based | Run with Docker container |

For each model and dataset, we used the area under the receiver operator characteristic curve (ROC AUC), using native-specificity variants as true positives and globally inactive variants as true negatives, as an initial metric of the model’s performance at their simplest, core task of identifying inactive variants. As expected, all tested models can separate inactive variants from native-specificity ones, albeit to varying degrees (Supplemental Figure 3). However, ProteinMPNN performs especially poorly for nearly all datasets; we thus excluded ProteinMPNN from our subsequent analysis. ESM-IF1 and the PSSM also performed slightly worse than the others at this task, which is again expected as ESM-IF1 is optimized for variant effect prediction and the PSSM is context-free. Otherwise, most models perform similarly, with inter-dataset variability far larger than average differences in model performance, which is consistent with larger-scale analyses such as ProteinGym^13^.

By contrast, there is strong variation between models in how they score altered-specificity variants relative to globally inactive or native-specificity variants. For example, on the PSD95-PDZ3 dataset, a site-independent model trained on a large MSA scores altered-specificity variants nearly identically to native-specificity ones, while the protein language model ESM-1v scores them far more similarly to inactive (destabilized) variants (Figure 2A). We analyzed this trend more systematically across the entire database by quantifying the average score of altered-specificity variants relative to inactive or native-specificity variants with an altered-specificity index (ASI; Figure 2B), defined as:

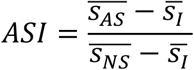

**Figure 2.**
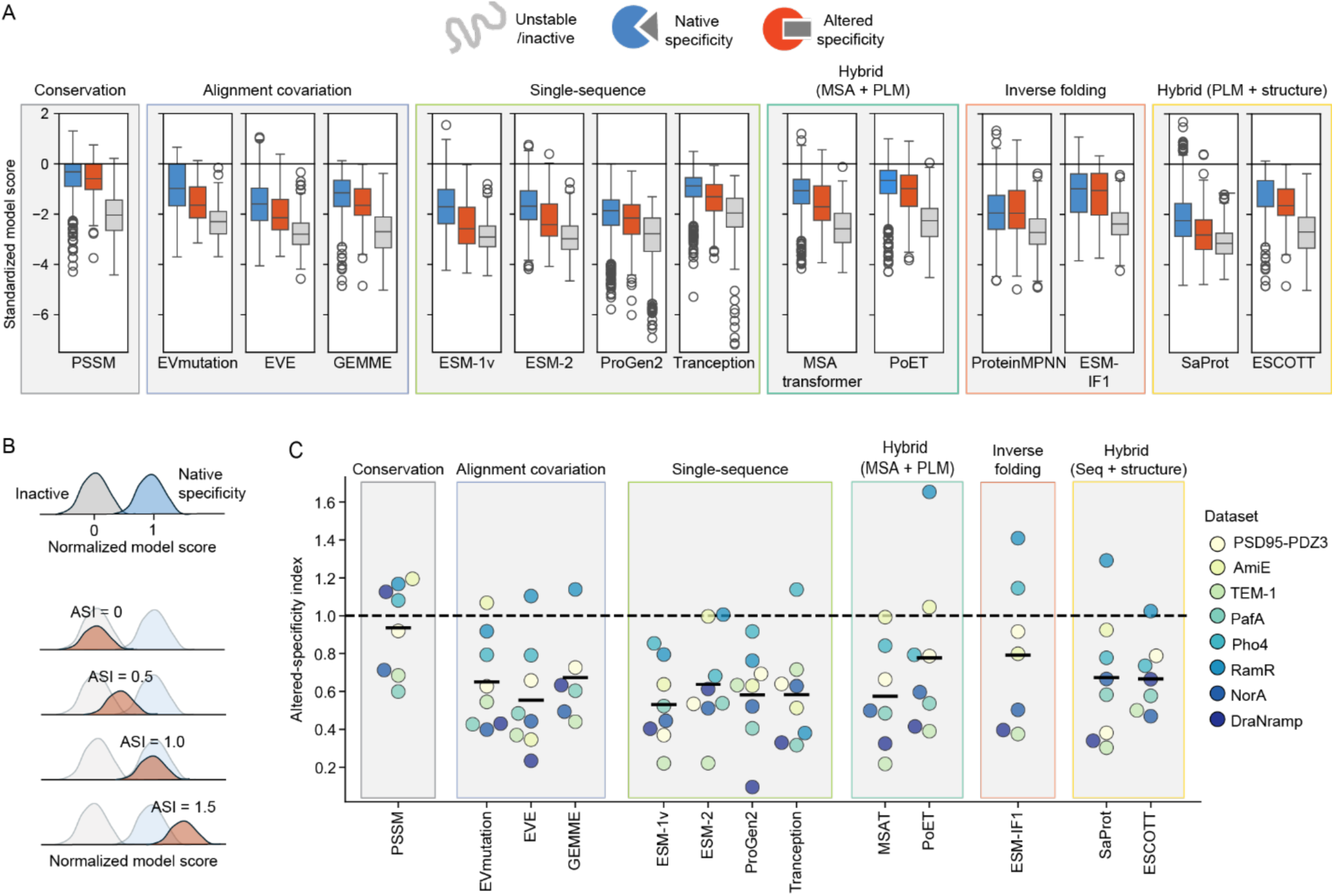
Zero-shot models that learn from sequence context systematically bias against altered-specificity variants. (A) Boxplots showing fitness scores (log-probabilities) assigned by each model to PSD95-PDZ3 variants separated into three classes: inactive (in this case, destabilized) in gray; native-specificity variants in blue; and altered-specificity variants in red. To place model scores on a comparable scale, we standardized them by dividing by their standard deviation and setting their score for the wildtype protein as zero. Models are separated by category, as defined in Supplemental Note 1. (B) Schematic illustration of the altered-specificity index (ASI). For each model, scores are first normalized such that native-specificity variants have a mean score of 1 and inactive variants have a score of 0. The ASI is the average score of altered-specificity variants on this scale. (C) ASI values for each dataset and model, with mean ASI values across datasets shown as a horizontal black line. Models are arranged as in panel A, except that ProteinMPNN was removed as it did not separate the native-specificity and inactive variants well enough to calculate ASI values. Individual points are colored by dataset.

Here, 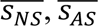, and 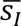 for each model are the average scores it assigns to native-specificity, altered-specificity, and inactive variants, respectively. An ASI of one indicates no systematic discrimination between native-specificity and altered-specificity variants, while an ASI of zero indicates that the model scores altered-specificity and fully inactive variants identically.

The protein language models and alignment covariation models have the lowest ASI values, indicating that they most strongly bias against altered-specificity variants (Figure 2C). Only the PSSM has a mean ASI near one, suggesting that it treats native- and altered-specificity equivalently. Hybrid models such as PoET, SaProt, and ESCOTT have intermediate ASI values, as do EVmutation and GEMME, which are able to condition on covariation information but in a less flexible way than deep-learning based models such as ESM-1v, EVE, or MSA transformer. Notably, the inverse folding model ESM-IF1 behaves most similarly to the PSSM, with relatively little bias between native- and altered-specificity variants. This is consistent with the growing consensus that log-likelihood scores extracted from inverse folding models report more closely on stability than on biochemical activity^13, 35, 36^, as altered-specificity variants are not necessarily less stable than native ones.

Comparison of AUC and ASI scores across datasets shows considerable heterogeneity in how models make predictions for each protein that likely relates to the nature of the biological questions (Figure 3). For the PSD95-PDZ3 dataset, there is no strong correlation between AUC and ASI (Figure 3A), and the PSSM separates native-specificity variants from inactive ones as well as ESM-1v or EVE. Previous studies have shown that EVE and ESM-1v both outperform a PSSM at predicting binding to the native CRIPT ligand^9, 13, 37^, and combined with our SpecificityStudio result this suggests that this performance gain is driven by their preferential prediction of native activity over altered activity. Doing so requires knowledge of what “kind” of PDZ sequence PSD95-PDZ3 is, which these models can do because they learn from context.

**Figure 3.**
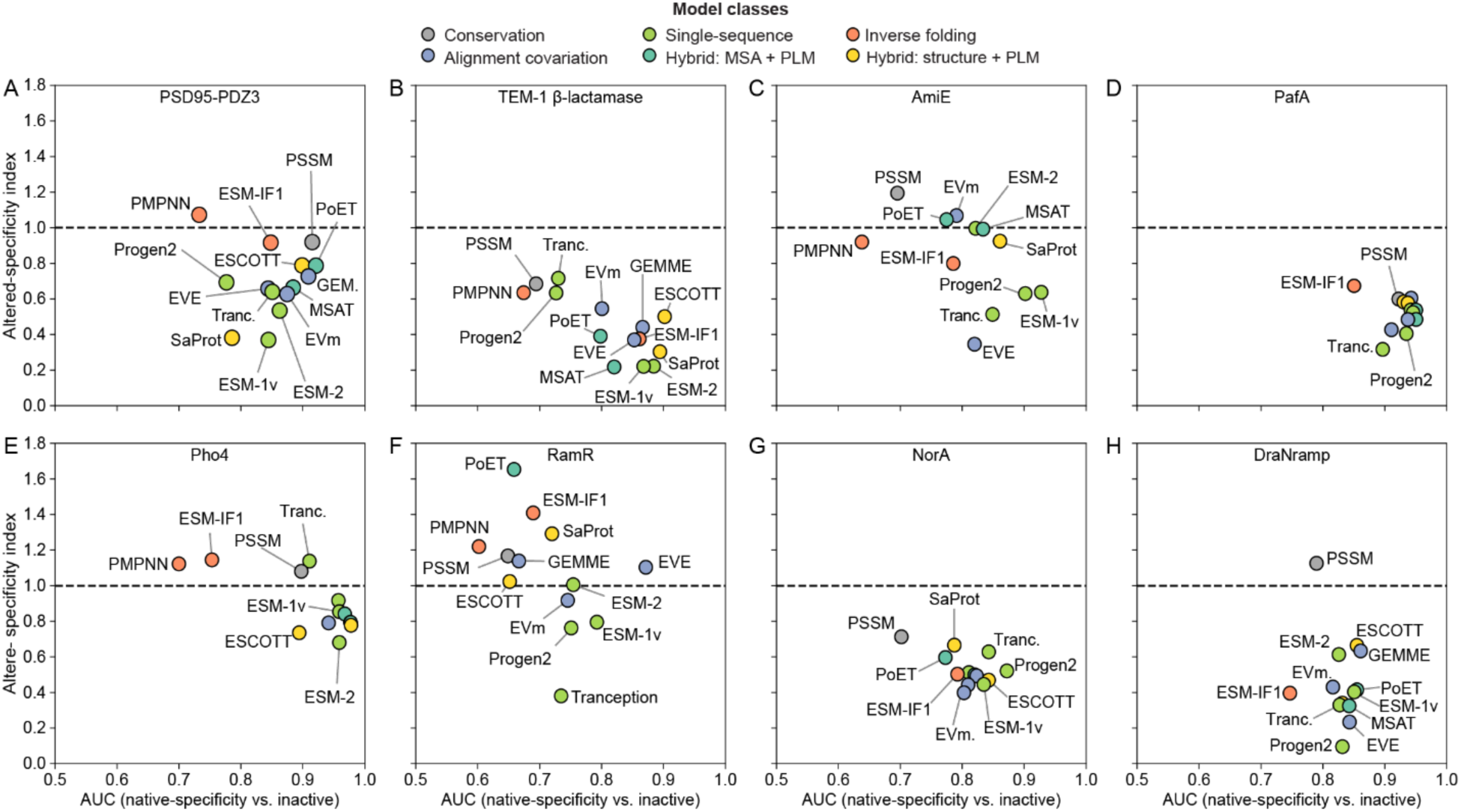
Trends in how models score different classes of mutations across datasets. Comparison of how each model separates native-specificity variants from inactive ones (“AUC”, x-axis) from how they score specificity-altering mutations relative to these (“Altered-specificity index”, y-axis) for each of the eight datasets. Models are colored by class and labeled when possible. Model names abbreviations: EVm; EVmutation; GEM. (in panel A), GEMME; Tranc., Tranception; MSAT, MSA transformer; PMPNN, ProteinMPNN.

A similar but more extreme trend can be observed for the DraNramp dataset (Figure 3H), where all models perform nearly identically at separating native-specificity from inactive variants but all models other than the PSSM have very low ASI values, with some nearing zero. Unlike many of the other datasets, DraNramp is under strong selection not to import Mg^2+^, and while there are likely many Mg^2+^-transporting variants in its alignment^38^, they have very low sequence identity. It is thus perhaps not surprising that learning from context especially biases against altered specificity in this case. The miocrofluidics-based datasets for PafA (Figure 3D) and Pho4 (Figure 3E) also show a similar trend but with relatively compressed values, perhaps influenced by the relative subtle substrate or ligand perturbations introduced in these experiments.

By contrast, the other datasets have narrower or less clear trends. For NorA, TEM-1, AmiE and RamR (Figure 3B, C, F, G), the PSSM biases less against altered specificity than most models but is also worse as distinguishing inactive variants, leading to an anticorrelation between AUC and ASI. Notably, the active/inactive labels in these cases are based on a relatively small set of substrates (3-11), unlike the near comprehensive substrate space explored in the PSD95-PDZ3 case, thus some mutations assigned to the inactive category may be functional on an untested axis. Alternatively, epistasis learned by the more complex models may contribute more to the predictions in these cases than for the simple binding domain PSD95-PDZ3, although the PSSM also performs very competitively for some complex proteins such as PafA and DraNramp.

Together, these results show considerable heterogeneity between datasets but nonetheless one clear trend emerges: protein fitness models that more strongly condition on sequence context—primarily variational autoencoders and protein language models—score mutations that alter protein substrate specificity much worse than those that do not perturb specificity. Only a site-independent model with no ability to learn from sequence context escapes this trend.

### Protein language models mostly design single variants with native specificity

The systematic bias against altered-specificity variants by models that learn from sequence context has consequences for what kinds of variants they design. To illustrate this, we simulated “designing” 5% of the total tested single-mutation PSD95-PDZ3 and NorA variants or double-mutation DraNramp variants (Figure 4A-C). We picked these three cases as representative examples as the available datasets: each have many mutations altering specificity, and these mutations do so in different ways. In each case, ESM-1v designs entirely avoid sampling variants with improved activity for an alternate ligand or substrate (PSD95-PDZ3 and DraNramp) or make the protein more selective for one of its native substrates (NorA).

**Figure 4.**
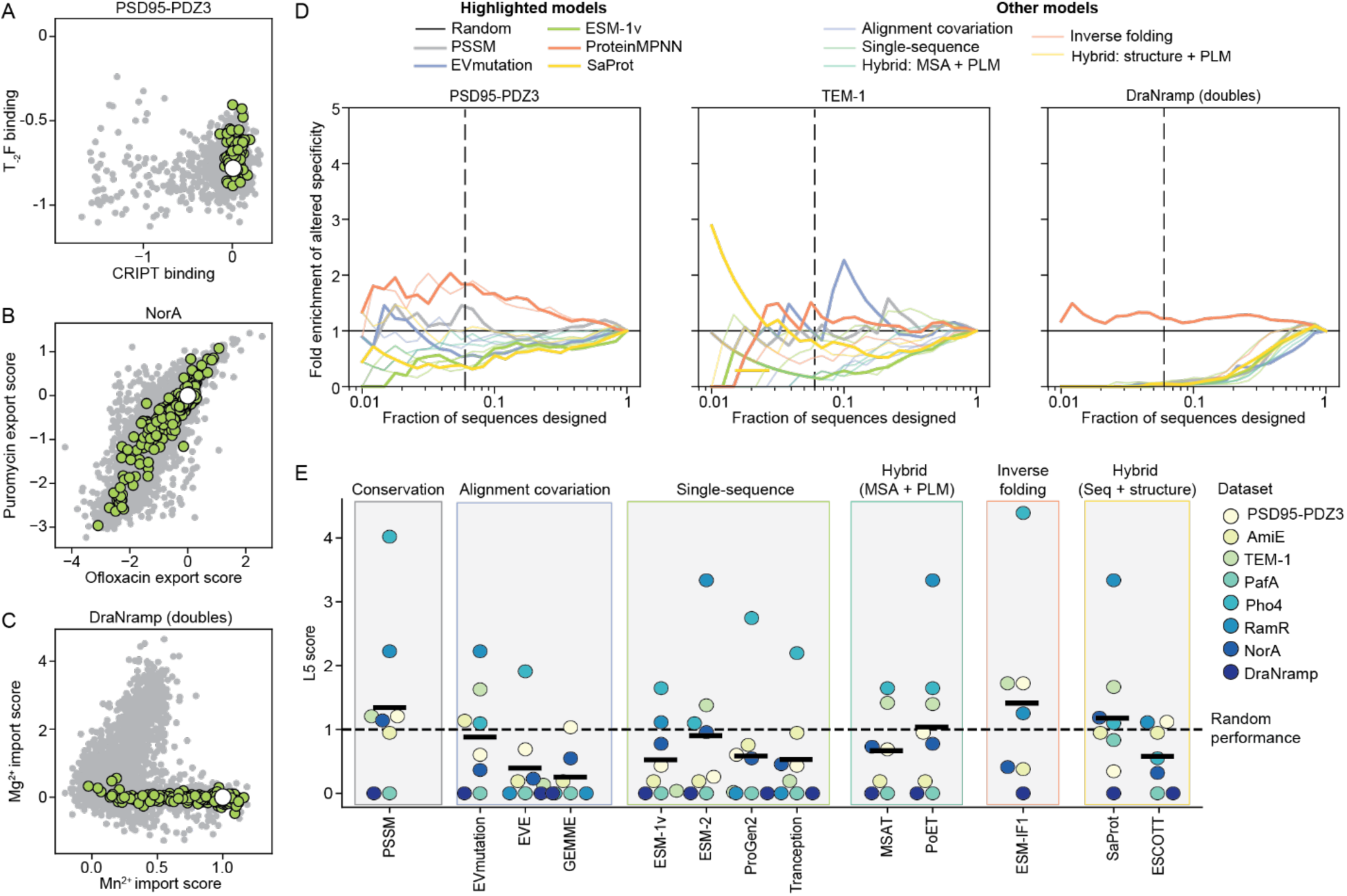
Zero-shot models that learn from sequence context generate mostly native-specificity variants. (A) Binding activity of PSD95-PDZ3 variants to its native ligand CRIPT and an example alternate ligand, T-2F, for the 250 (or 5%) top simulated single-mutation “designs” from ESM-1v (green). All variants in the dataset are shown in gray. All samples from ESM-1v bind CRIPT well, but none switch its specificity toward T-2F. (B) Similar simulated design analysis for NorA, where single-mutation designs from ESM-1v tend to perturb ofloxacin and puromycin export similarly, almost none differentially affect transport of these two substrates. (C) For double mutations to DraNramp, ESM samples no variant that transports Mg^2+^ in the designs scoring in the top 5%. (D) Examples of altered-specificity enrichment curves for single-mutation variants of PSD95-PDZ3 and TEM-1 or double-mutation variants of DraNramp. Each curve represents the enrichment of specificity-altering mutations within a fixed budget of N sequences designed by the model in question. A vertical line at 5% represents the fixed cutoff used as a summary statistic. (E) The enrichment of altered-specificity variants when designing 5% of the total sequence library for each dataset (the “lift at 5%” or L5 score) and each model, with the mean shown as a black line. All datasets are limited to single variants. Most models have average performances at or below random chance.

We quantified the utility of each model for designing variants of new specificity by generating an altered-specificity enrichment curve. We simulated “designing” the top *n^th^* percentile sequences from each model across all datasets and calculated the enrichment of altered-specificity variants in the designed set compared to random chance. This analysis quantifies the increase of performance of model-guided design across different “budgets” of designed sequences compared to random chance (Figure 4D and Supplemental Figure 4). The broad utility of sequence data for this task varies considerably by dataset, as evidenced by the very different performances of the PSSM (gray in Figure 4E). In some cases, site-independent evolutionary information leads to >4x enrichments of altered-specificity variants (e.g. for the RamR dataset) while in other cases this information leads to sampling altered-specificity variants at a rate far below random chance (e.g. the DraNramp dataset; Figure 4E). Averaged across all datasets, using site-independent evolutionary information is approximately neutral for design success across the different datasets, consistent with the idea that this information is generally impartial on the matter of altered versus native specificity. The inverse folding models are slightly better at enriching libraries, although once again their performance is highly variable and difficult to distinguish from random chance.

Most concerning are the performances of protein language models and covariation-based alignment models. Even though these models are excellent at separating native-specifity active from generally inactive variants (Supplemental Figure 3), their biases against altered-specificity variants mean that they design fewer such variants than random chance, often significantly so. On several datasets, ESM-1v, Progen2, and Tranception each design no altered-specificity variants in the 5% top-scoring designs tested, nor do EVE, GEMME, or ESCOTT. These results suggest that using these models to guide initial exploration in a directed evolution campaign for altered function would likely have poor results.

### Differences between protein fitness models can help prioritize variants likely to alter specificity

No tested model on its own has even a two-fold average enrichment of altered-specificity variants over random chance (Figure 4E). However, the large differences between scores assigned by different models suggest that models learning from sequence context must implicitly encode information related to specificity and this information could still be used to guide library designs. As an analogy, a student who gets every question wrong on a multiple-choice exam must have known some of the answers.

Due to the limited size of our dataset, we took a conservative first approach to learning from model differences by fitting weighted ensembles of pairs of models. We performed logistic regression against variants classified as having altered specificity across datasets to fit a single global parameter, α, weighing the two predictors (Figure 5A; see *Methods*). In addition, we fit dataset-specific logistic regression parameters to account for different thresholds of specificity alteration per dataset. To evaluate the dataset-specific predictive performance, we alternately held out each dataset from the training to fit α and then assessed its design enrichment curve with the weighted predictor as the protein design model.

**Figure 5.**
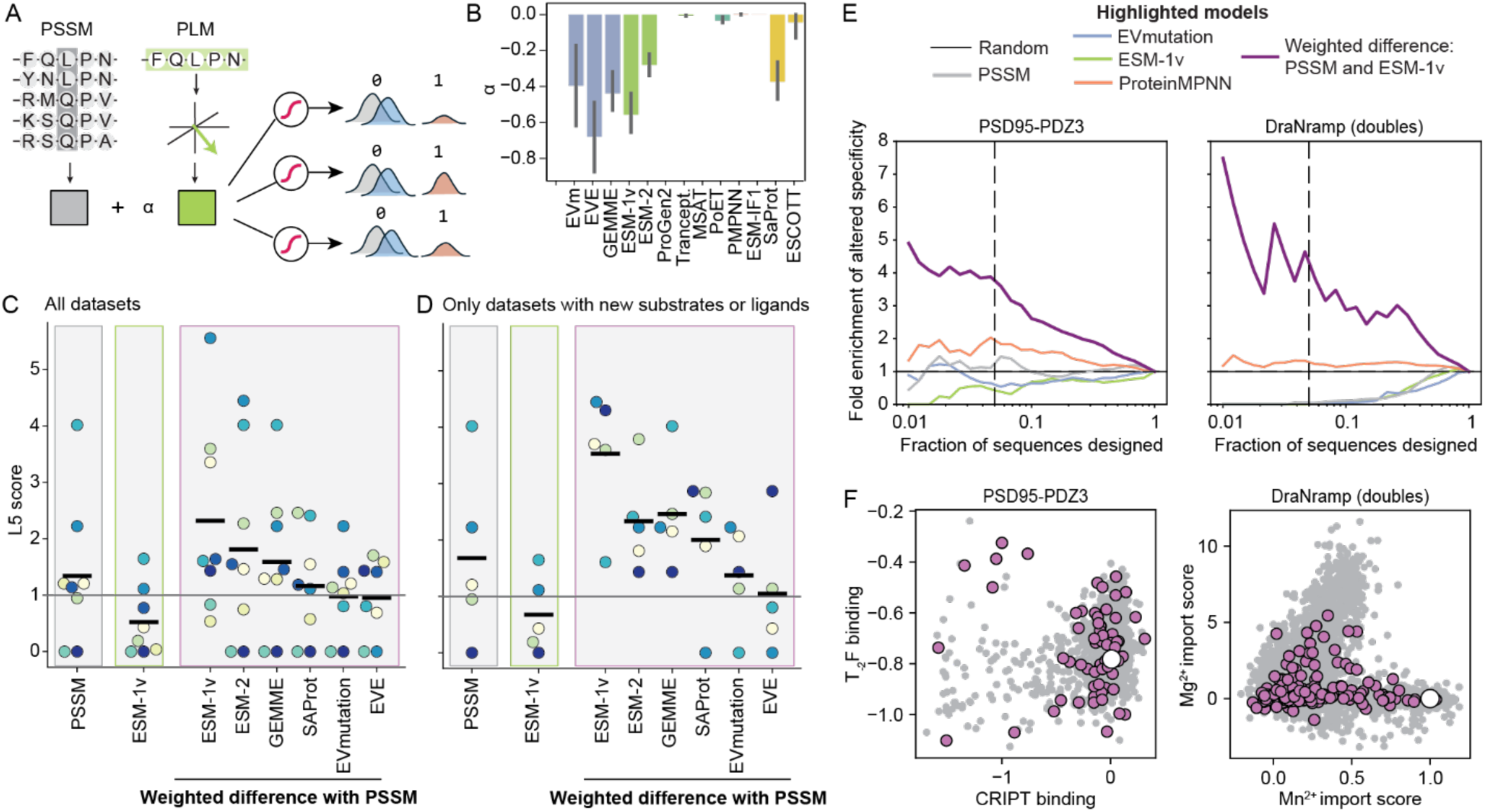
Weighted differences between models can enrich design libraries for altered-specificity variants. (A) Schematic of the weighted difference model inference, showing how we learn an optimal weighting between a site-independent model (PSSM) and model that learns from context such as a protein language model (PLM) by performing logistic regression against each dataset (see *Methods* for details). (B) Optimal α values fit via holding out each dataset for ensembling each model with the PSSM across all datasets. All weights are either zero (suggesting the additional information is not useful) or negative (suggesting that a weighted difference helps discriminate altered-specificity variants best); none fit a positive-weighted ensemble. Error bars represent a 95% confidence interval across training sets. (C) The enrichment of altered-specificity variants when designing the 5% of tested variants ranked highest by each model, called the “lift at 5%” or L5 score. The PSSM and ESM-1v are included on the left (reproduced from Figure 4E), while different weighted ensembles with the PSSM as the first model are shown in purple and ordered by mean enrichment. When fitting an optimal weighting for each model and dataset, that dataset is held out from the training data, leading to a slightly different α value for each dataset. Individual scatter points are colored according to dataset as in Figures 2 and 4. (D) The same type of plot as panel C but with only Class 1 datasets. Most models perform slightly better, and a weighted difference with ESM-1v leads to 4-fold average enrichment. (E) Example specificity enrichment curves (as in Figure 4D) showing the improved performance of a weighted difference between the PSSM and ESM-1v in purple. (F) Similar plots to Figures 4A and 4C but “designing” the top 5% of labeled sequences from PSSM-ESM-1v weighted difference model. Many sampled sequences have altered specificity, either by switching PSD95-PDZ3 specificity to T_-2_F or allowing for Mg^2+^ import for DraNramp.

Every model when ensembled with the PSSM either has a negative or zero optimal weighting (Figure 5B), suggesting—as we hypothesized—that a weighted *difference* is the most effective approach rather than an additive ensemble. A weighted difference between the PSSM and ESM-1v led to the greatest enrichment, although only approximately 2.5-fold (Figure 5C). We reasoned, however, that one complicating factor leading to highly divergent behavior between assays might stem from the different kinds of specificity alteration present in each dataset. We therefore split the assays into two categories, the datasets representing improved activity against ligands or substrates that are non-native to the wildtype protein (Class 1: PSD95^pdz3^, TEM-1, Pho4, RamR, DraNramp) and those representing modulations in the relative specificity for different native substrates (Class 2: AmiE, PafA, and NorA; see Supplemental Note 2 for reasoning for each dataset). Despite a reduction in training data, fitting weighted ensembles only on Class 1 datasets improved the average enrichment of altered-specificity variants among the top 5% of designs to 4-fold for each held-out Class 1 dataset (Figure 5D-F). The largest increase in performance was for the DraNramp dataset (darkest blue in Figure 5D), for which it is advantageous to strongly weigh the negative term for ESM-1v predictions. This is notable as DraNramp is the only dataset for which the wildtype has truly no measurable activity on the non-native substrate. Together, these results suggest that differences between context-independent and context-aware models are especially useful for designing major shifts in specificity away from the wildtype protein’s ligands or substrates, rather than subtle modulations to the specificity profile for existing substrates.

## Discussion

### Learning from sequence context biases models against altered specificity

In this study, we built the first dedicated database of diverse multiplexed functional assays against multiple substrates and identified from these datasets which variants alter each protein’s substrate or ligand specificity. We then analyzed the fitness (log-likelihood) scores that 14 different zero-shot sequence- or structure-based models assign to these variants and identified a systematic bias against altered-specificity variants in context-aware models. Because this bias leads to undersampling altered-specificity sequences compared to random chance, it can be exploited by training weighted ensembles that take the difference between scores from different models to enrich simulated library designs for altered-specificity variants.

The strong bias against altered-specificity variants by context-aware models makes sense based on how these models make a prediction. As a toy example, we consider a family of proteins with two distinct substrate specificities (blue or green in Figure 6). Different mutations at the same position could maintain specificity (mutation to an amino acid associated with substrate 1) or alter specificity (mutation to an amino acid associated with substrate 2). For a context-free model that simply uses the amino acids frequencies in each column independently, both amino acids are present in the natural sequences and thus they are assigned approximately equal probabilities (Figure 6B). However, models such as EVE and ESM-1v learn from the entirety of the sequence to generate some form of latent representation (*z* in Figure 6C), These latent representations guide such a context-aware model to learn more from more similar sequences and thus assign higher probabilities to the amino acids found at that position in more similar sequences.

**Figure 6.**
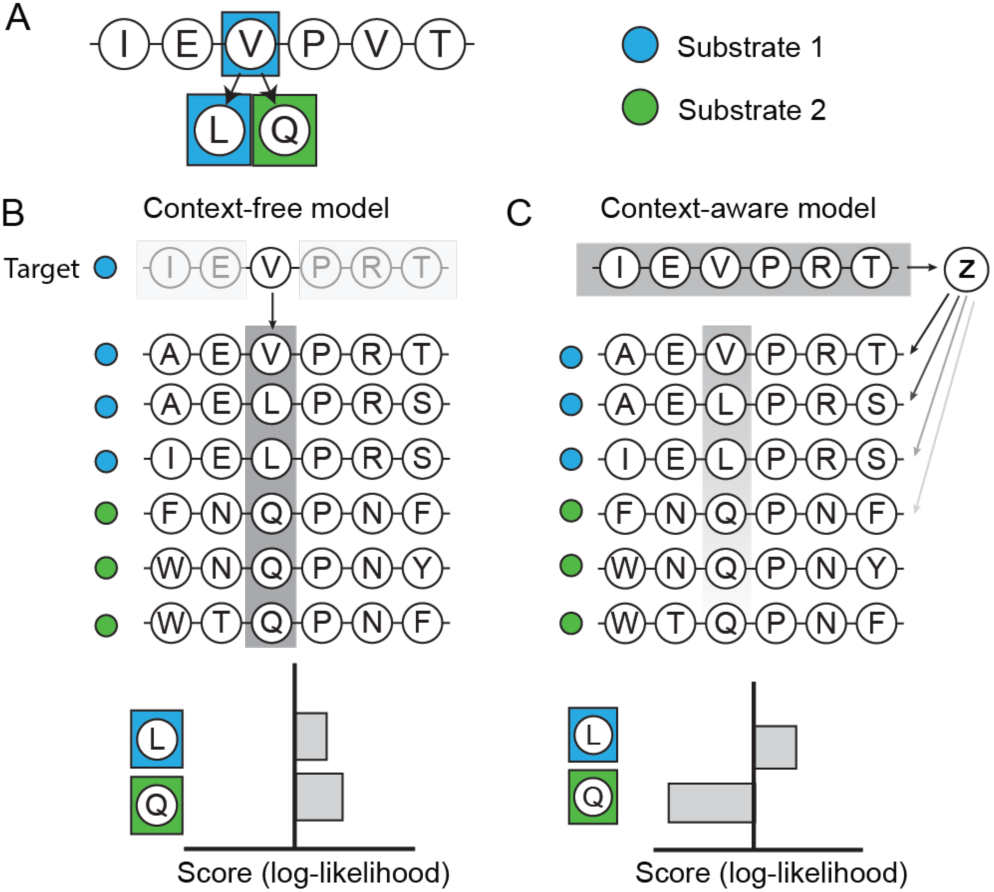
How context-dependence can shape fitness models’ predictions for altered-specificity variants. (A) Illustration of a sample protein variant effect prediction task, where a model is being asked to choose between two mutations: V→L which preserves the protein’s native specificity for its substrate 1 (blue), or V→Q which alters the specificity to substrate 2 (green). (B) Illustration of what a context-free model might learn from an MSA of homologs. Colored circles on the left represent the true specificity of each natural sequence. Because the model assigns its scores purely based on one position at a time, without learning from sequence context, it simply sees a similar number of leucines and glutamines in the MSA column and assigns them similar, positive scores. (C) Illustration of a context-aware model such as EVE or ESM-1v. In this case, the model uses the entire sequence to embed category information for each target protein into some additional parameter z. In a prediction, z leads the model to focus its search primarily on sequences more like the target (those with a similar specificity), and thus it assigns a far higher score to the leucine mutation than to the glutamine mutation.

While a relatively straightforward consequence of how these models make predictions, the finding that EVmutation, EVE, and – especially – ESM-1v bias against altered specificity goes against an assumption made by many recent studies that these models will be useful for engineering altered functions because they learn a generalized notion of protein fitness^21–24, 39–41^. In some cases, the functions tested are enhancements to the existing, evolved properties of the protein^24^. However, some studies did successfully use ESM log-likelihoods to design altered function but in the context of antibody-antigen binding^22, 40^. In antibody-antigen binding the evolutionary sampling process is far more random and less contingent than the slow, stepwise natural evolution of typical protein domains. We chose to exclude antibodies from this analysis for this reason and because of the difficulty of generating antibody alignments, but a systematic comparison of predicting specificity for antibodies compared to other protein families would shine light on these distinctions.

Still, our demonstration of bias against altered-specificity variants across diverse types of proteins shows that context-aware models such as protein language models do not learn general evolutionary rules for most protein families. Instead, it suggests that protein language models learn implicit mixtures of function-specific modes that exclude regions of sequence-function space that have not been sampled by natural evolution. These results should hopefully serve as a reminder and word of caution to researchers interested in using protein language models for engineering. Unsupervised sequence models have learned to do exactly what they are trained to do: to very closely fit the *observed* probability distribution over natural sequences in databases, which represents a nonequilibrium sampling from the true space of fitness landscapes. Only models trained using different objectives (e.g. inverse folding models) or unable to learn from sequence context (conservation models) avoid closely fitting these observed probability distributions closely and thus minimize these biases. In a related context, this principle has been referred to as the “blessing of misspecification” for models of molecular fitness^42^.

### Recommendations for protein engineers

Our results emphasize the need for caution when applying zero-shot predictions to protein engineering. Models that predict native function well – for example, due to high scores on ProteinGym – do not necessarily generalize to predict altered function, and indeed the correlation is often inverse. When choosing a model trained on evolutionary data, a site-independent model is more likely to be able to generalize to an alternate function than a protein language model, which would be more useful for redesigning a protein sequence to enhance a function that was already actively under selection. This is also likely highly dependent on what sequences are in the alignment. We focused primarily on very large sequence alignments that contain within them a wide array of specificities; for a protein family for which this was not possible, alignment-based information is likely less useful.

Even still, no zero-shot method on its own is very useful and it is almost certainly more practical to simply test mutations at random. Only by learning from differences between model predictions were we able to guide simulated libraries that significantly outperform random chance. In particular, taking a weighted difference between a site-independent PSSM model and ESM-1v is likely to enrich for variants that alter specificity in new datasets. We provide code for scoring sequences and sampling directly from such an ensemble given an input MSA.

### Limitations and future directions

The weighted difference approaches presented above can enrich library designs several-fold above random chance, but we do not mean to suggest that these models represent the most that can be learned from natural sequence data. They more likely represent a floor to how these data could be used with additional experimental benchmarking and principled modifications to the language models themselves.

The primary limitation of our study is the small size and heterogeneous nature of the datasets in SpecificityStudio. This is largely because most mutational scanning studies to date have focused on a single substrate or ligand at a time. However, recent advances in the scaling of highly quantitative, massive-scale measurements of protein function^43, 44^ make it likely that such data will become available soon. For instance, while our assembled datasets include one binding domain-peptide pair with data for all-against-all single variations, large-scale assays could profile hundreds or thousands of different binding domains against diverse arrays of substrates. Indeed, two studies moving in this direction were released while this manuscript was in preparation^45, 46^. A similar analysis could be done for transcription factor binding, allosteric transcription factor activation, enzymatic catalysis, or any other class of protein activity. It is our hope that a future update to SpecificityStudio may contain such data.

In the meantime, and as new datasets are released, it would be prudent for developers of machine learning models to keep in mind not simply how their models score effects on a protein’s native function but how they score effects on alternate functions such as interactions with non-native ligands or substrates. This might involve analyzing a model’s bias against altered-specificity variants but also developing ways to control how a model samples sequences with native-like specificity versus ones that remain functional on some axis but deviate more from the native-function of its nearest evolutionary neighbors.

In this study, we focused exclusively on zero-shot predictors that condition only on sequence and possibly a structural model. However, many computational models can now make predictions conditioning both on natural sequence information and on the actual ligand or substrate tested^39, 47–49^, information that we did not use in this study. Information about the ligand or substrate in question is clearly necessary to go beyond simply enriching libraries for generally active or even generally function-changing sequences and make actual predictions about specific activities. Models such as Boltz2^47^ have tested their ability to predict binding free energies based on datasets of protein-small molecule binding free energies, but testing their ability to predict perturbations in binding affinity upon mutation (as in SpecificityStudio) would provide an orthogonal and stringent test of how well they have learned the principles of binding. However, only two datasets in SpecificityStudio report on binding directly, and it remains to be seen whether any model that conditions on static structures would be useful in predicting interactions with different molecules when the downstream phenotype also requires protein dynamics, particularly for the transporter and allosteric transcription factor datasets.

We also did not touch on the broader question of different models’ utility in semi-supervised prediction. Semi-supervised models trained both on natural sequence information and a limited set of experimental labels have shown improved performance over zero-shot models on their own^50–52^, although they have generally been evaluated on mutational scanning studies with a single functional readout. It is unclear how semi-supervised models trained on foundations that bias against specificity-altering mutations will perform at predicting function with different substrates or ligands given limited experimental data. While a small amount of training data may guide a model to make different predictions, the biases stemming from the unsupervised portion of the model may lead to difficulties learning the non-native landscape. We hope to see the datasets in this and future versions of SpecificityStudio used for these benchmarking tasks and others we have not yet imagined.

## Methods

### Building MSAs

All MSAs were built using JackHMMER^53^ as implemented in the EVcouplings pipeline^54^, except for the Pho4 alignment. Due to the small size and high sequence diversity of the bHLH transcription factor family, we were unable to generate a reasonable alignment with JackHMMER and thus followed Hastings et al^55^ in using a pre-curated alignment from InterPro representing the entire bHLH family. To match this alignment to the full Pho4 sequence, we then fetched all of the full-length original sequences from UniProt with easel-sfetch^53^ and realigned them to a seed HMM built from the original alignment, then filtered to columns present in yeast Pho4. Otherwise, we ran JackHMMER for five iterations with a variety of relative bitscore thresholds and picked the most diverse alignment that we could generate. Table 3 summarizes the alignments used. The effective number of sequences is calculated as in EVcouplings^54^ with θ = 0.8.

**Table 3.**
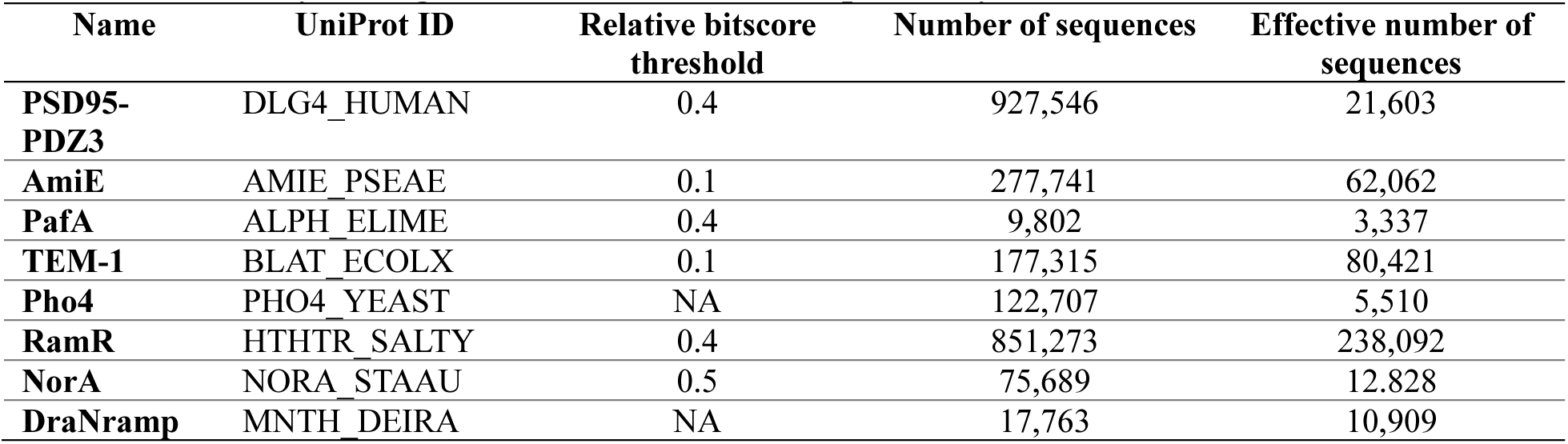
Summary of alignments for datasets in SpecificityStudio.

### Scoring mutation effects

Fitness model scores (log-likelihoods) were calculated using scripts directly from ProteinGym^13^ with several exceptions. GEMME^56^ and ESCOTT^57^ have complex dependencies that are difficult to run on computing clusters; GEMME and ESCOTT were therefore both run using their provided Docker containers. EVmutation^54^ and site-independent scores were additionally calculated with a custom script.

### Identifying altered-specificity variants with linear regression

For the PafA and Pho4 datasets we used a linear regression approach to identify altered-specificity variants. In each case, we defined a “native” substrate/ligand baseline (MeP, 5’-CCACGTGA-3’, or 7-iodo-indole, respectively) and performed least-squares linear regression against scores with the “native” substrate/ligand and each other substrate/ligand with scipy^58^. For PafA, the trend between MeP and cMUP catalytic efficiencies falls off for very low values of each. Therefore, we first filtered the dataset to include only variants with a k_cat_/K_M_ of 5000 M^-1^ s^-1^ for MeP and 20,000 M^-1^ s^-1^ for cMUP. We then separate each variant’s score on each substrate/ligand into its global and specific component by drawing a perpendicular line from it to the regression line: thus, the “global” activity value for a point with values (*x*_1_, *y*_1_) given a regression line with slope *m* and intercept *b* is given by:

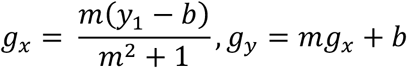

The specific activity is the distance from the original point to the intersection point, or:

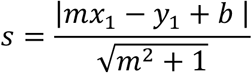

### Altered-specificity enrichment curve

For a given percentage of the data *p*, in a dataset with *N* total labeled variants, of which *k* variants alter the specificity, we take the *Np*-top scoring variants from a given model and define the enrichment as the number of altered-specificity variants in that set divided by the expectation from random chance 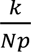. We then vary *p* on a logarithmic scale from 0.01 to 1. The L5 score represents the value of this curve when *p*=0.05 (5% of the total data).

### Fitting weighted difference models

We sought to use the data to infer a pair of models that would help enrich libraries most for altered-specificity variants. In general, such a metapredictor would simply define a new combined score based on the predicted fitness values of the two models:

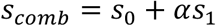

For each dataset *d*, we modeled the probability of a variant altering the specificity using logistic regression:

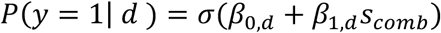

Here, σ is the logistic function and β_0,*d*_ and β_1,*d*_ are a dataset-specific bias and slope, respectively. The optimization then proceeds in two nested levels. First, for a fixed value of α, we fit the dataset-specific parameters with the weighted log-likelihood:

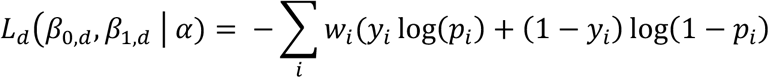

Where *w_i_* are sample weights. These weights allow for class balancing by weighing each sample inversely proportion to its class frequency. To optimize α, we then fit:

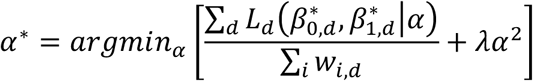

Where 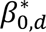 and 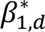 are the optimal parameters inferred during the inner optimization, and λ is a regularization coefficient. Note that normalizing the log-likelihood of each dataset by its sample weight ∑*_i_ w_i,d_* ensures that each dataset contributes equally to the overall objective function, even though they are different sizes. We perform the inner optimization (of the logistic regression parameters) with BFGS optimization and the outer optimization (of α) with L-BFGS-B bounded between -10 and 10. We use weak L_2_-regularization (λ=0.1) to prevent extreme values.

For every prediction in Figure 4, we excluded the dataset being evaluated from the training set, thus training a different set of α parameters for each dataset. When applied to a new dataset, this model cannot assign exact probabilities of specificity-switching, which requires fit β_0,*d*_ and β_1,*d*_ parameters to learn the exact cutoff that constitutes altered specificity. However, it can still be used to rank sequences, which we evaluate using altered-specificity enrichment curves and the L5 score.

## Acknowledgements

We thank members of the Marks and Gaudet labs for useful discussions. We would also like to thank Emma Garcia for general advice and feedback. This work was funded in part by NIH grants R01GM120996 and R35GM161633 (R.G.), the NSF-Simons Center for Mathematical and Statistical Analysis of Biology at Harvard (award number 1764269) and the Harvard Quantitative Biology Initiative (S.P.B.), NIH/NIGMS T32GM0008313 (S.P.B., PI Venkatesh N. Murthy), and the Chan Zuckerberg Initiative grant R01CA260415 (D.S.M.).

## Author Contributions

S.P.B. and D.S.M. conceptualized the project. S.P.B. ran models, performed all analyses, and wrote the manuscript with input from R.G. All authors reviewed and edited the manuscript.

