## Supplemental information (Supplemental Notes 1-2 and supplemental figures) for "Differences between protein fitness models can be used to design variants of altered specificity"

This file contains:

- Supplemental Notes 1-2
- Supplemental Figures 1-4

**Supplemental Note 1. Description of model classes**

To allow for a more in-depth analysis of results we focused on a smaller representative set of popular models in this study than in ProteinGym. We distribute these models into five classes: conservation, alignment covariation, pretrained single-sequence, structure-sequence ("inverse folding"), and hybrid models. We consider all models as probabilistic sequence models—that is, they attempt to fit the unknown "true" fitness distribution $p(x_i)$ for a sequence $x_i$ based on the observed sequences in either a multiple sequence alignment (MSA) or the entire universe of known protein sequences.

*Conservation models*

The simplest class of protein fitness predictors relies on residue conservation,[1,2] sometimes called site-independent predictors,[3] profile models,[4,5] or position-specific scoring matrices (PSSMs).[5] These models rely on an MSA of homologous sequences and implicitly infer the probability of each amino acid at each site based on its frequency in the alignment, often with some statistical adjustments depending on the exact algorithm.[4] A somewhat more sophisticated version of this algorithm that does not require alignment is called a *profile hidden Markov model* (HMM).[4] In our analysis, we used the site-independent PSSM model as a representative conservation model (see Table 2).

The actual implementation of conservation models is typically trivial given an MSA, and the key factor in prediction is the nature of the MSA used. While an exhaustive review of all kinds of alignment variability is outside the scope of this study, for each protein investigated, we build several different MSAs of varying depths, always using the Jackhmmer-based[6] protocol introduced in EVmutation.[3]

*Alignment covariation models*

In general, we classify all models that use an MSA but condition a variant's score on the amino acid identities at other positions in any manner as *alignment covariation models*. We distinguish these from position-independent alignment models ("conservation-based") as, rather than conditioning a given mutation effect on a set of terms specific to just that position, it conditions the effect on the entire sequence. Conditioning on the rest of the sequence can lead to very different behavior. The first of these models to be popularly used was direct couplings analysis (DCA),[3, 7-9] which we implement via the EVcouplings pipeline[10] and refer to as EVmutation when predicting mutation effects. EVmutation is an energy-based model that scores a mutation based on an explicit set of sitewise conservation terms and pairwise energetic coupling terms. These couplings, the strongest of which frequently predict structural contacts,[7, 11] are often interpreted as representing epistatic interactions.[3, 9] Due to the non-identifiability of fitness and phylogeny,[12] however, the covariation underlying these couplings could equivalently arise from phylogenetic or functional structure in the MSA. Other models, such as the variational autoencoders (VAEs) DeepSequence[13] and the EVE[14] model included in our analysis, explicitly condition on the sequences' representation in a low-dimensional learned underlying sequence space. This sequence space naturally captures variability in the MSA and conditions the models' prediction based on it, although this relationship is also often described as indirectly encoding epistatic relationships.[13] Some models, such as GEMME,[15] accomplish a similar conditioning on phylogenetic or functional relationships but use an ensemble of predicted trees without any deep learning. The authors of this study also describe its predictions which incorporate phylogenetic information as "epistatic."[15] GEMME, which we included in our analysis, is notable for its strong performance on the

ProteinGym benchmark set[16] despite its relative simplicity and extremely fast run time compared to other methods.

*Single-sequence models*

Single-sequence models such as protein language models (PLMs) do not explicitly leverage an MSA but instead pretrain on all proteins in a large sequence dataset. In the case of PLMs and related models, a variant effect is predicted by conditioning on a very high-representational embedding of the sequence context. The most widely PLMs used today are the evolutionary-scale model (ESM) family. The first-generation ESM was trained as a masked language model with a deep bidirectional transformer architecture akin to BERT[17] with two similar flavors for protein variant effect prediction, ESM-1b[18] and ESM-1v.[19] Each flavor of ESM-1 predicts the effect of mutations based on the log-odds ratio at the mutated position, assuming no epistasis and thus combining multiple mutations additively. ESM-1v, which we included in our analysis, is trained on Uniref90 and typically runs as an ensemble of five models, each of which underperforms ESM-1b—trained on UniProtKB—but which together have been suggested to slightly outperform it, although the differences between ESM-1b and ESM-1v are likely very minor. Shortly after, the team also released ESM-2 (included in our analysis), updated with more parameters and optimized to predict structure,[20] although it can—and has been[21, 22]—equivalently used for variant effect prediction and design.

Many other protein language models have been developed and deployed for various uses. These include a family of autoregressive language models, ProGen,[23] ProGen2,[24] and ProGen3,[25] which have been experimentally demonstrated to generate functional sequences and used to engineer new CRISPR-Cas9 proteins.[26] Tranception[27] is another transformer-based autoregressive language model with similar performance but with the ability to integrate inference-time retrieval (see *Hybrid models* below). We included ProGen2 and Tranception in our analysis.

*Structure-sequence (inverse folding) models*

In contrast to the purely sequence-based models described above, so-called inverse folding models—beginning with ProteinMPNN[28, 29] and then ESM-IF1[30], both of which are included in our analysis—are trained to learn the probability of a sequence given a structure.[31] As such, rather than simply training to match the protein sequence distribution in a large database, ProteinMPNN was initially trained to learn the matching between all pairs of structures and sequences in the PDB,[28] while ESM-IF1 was additionally trained against all predicted structures in the ESMFold database.[30] Although these models were initially intended for design of full sequences from structures, their log-likelihood scores can also be used to score individual protein variants[30] and correlate reasonably well with stability but less well with protein activity or broader fitness.[16] This matches an expectation that these models learn a probability distribution that is more closely related to the protein folding landscape than the broader protein function landscape. It has been suggested that this can be used to discriminate between deleterious variants that decrease stability from those that break other functions[32].

*Hybrid models*

Many newly developed machine learning models use some combination of pretraining on the protein universe, retrieval of a particular MSA, and incorporating information from protein structures. Models that pretrain on all sequences but additionally use MSA information include MSA transformer,[33] Tranception with retrieval,[27] TranceptEVE[34], and PoET.[35] Some of these models—especially TranceptEVE and PoET, the two we included in our analysis—have consistently scored slightly better on the ProteinGym benchmark than purely alignment-based or purely single-sequence models. Among these, PoET (which stands for Protein Evolutionary Transformer) performs especially well in ProteinGym and has a unique architecture as an autoregressive language model that formulates a set of homologous sequences as a "sequence-of-sequences."[35] Other models that combine single-sequence information with structural context include SAProt,[36] ProSST, and S3F, while models that combine MSAs with structural information include ESCOTT[37], VenusREM[38], and S3F-MSA[39]. (SAProt and ESCOTT are included in our analysis.) All these models but ESCOTT use transformer-based deep learning architectures, while ESCOTT—developed by the same group that developed GEMME—adapts GEMME's phylogenetic-based approach while integrating structural tokens. Much like GEMME, ESCOTT matches the state-of-the-art performance on ProteinGym and even surpasses many models with far more parameters that are vastly more expensive to train. The popular but closed-source model AlphaMissense[40] also uses both sequence and structural information but is excluded for all analyses as scores are not available. The subset of mutational scanning predictions that are publicly released for AlphaMissense suggests that it is a strongly performing model, but—unlike AlphaFold, which is consistently better than its competitors—it is relatively standard among the latest generation of models integrating structural and evolutionary information.

**Supplemental Note 2. Description of mutational scanning datasets**

Below we briefly describe each dataset included in SpecificityStudio (see Table 1), how we inferred altered-specificity variants, and categorized each dataset into one of two classes: Class 1 datasets profile shifts toward new substrates or ligands, and Class 2 datasets profile shifts in specificity among existing substrates or ligands of the protein.

*PDZ domain PSD95-PDZ3*

PDZ domains are one of the largest families of modular protein binding domains with hundreds of members in most proteomes[41]. Most PDZ domains recognize the extreme C-terminal region of various proteins, with a wide array of specificities, to drive diverse protein-protein interactions. PDZs are broadly sorted into specificity "classes" mediated most significantly by the amino acid identity at the -2 position of the recognized C-terminal peptide,[41] although the true domain—peptide specificity mapping is complicated and, despite years of research, only partially understood.[42, 43]

The third PDZ domain of PSD95 and its cognate peptide of CRIPT[44] have long served as a model for PDZ function and for protein binding in general. A seminal mutational scanning study in 2012 assessed the binding of every single variant of $PSD95^{pdz3}$ to CRIPT and a "class-switched" single mutant to CRIPT, $T_{-2}F$, using a bacterial two-hybrid assay.[45] A more recent expanded dataset reports on every single variant of $PSD95^{pdz3}$ against every single variant of CRIPT using a related BindingPCA assay in yeast.[46] A similar AbundancePCA assay has also been used to profile the abundance of every single mutant and many multiple mutants to $PSD95^{pdz3}$,[47] serving as a useful complement to the binding data. For each of these studies, the authors performed 3-5 replicates, which correlate exceptionally well. Together, these datasets paint one of the most systematic portraits to date of the specificity design rules in a protein-ligand binding, making it an exceptional dataset for evaluating the behavior of various classes of computational models.

To define altered-specificity variants in PSD95$^{pdz3}$, we chose to directly use Zarin and Lehner's FDR-based metric[46]. Briefly, they began by considering the domain and peptide as a single folding reaction and thus proposed the null hypothesis that mutation effects on the binding energy at the domain level and mutation effects on the binding energy at the peptide level should be additive. They then fit a thermodynamic MoCHI model that accounts for global epistasis by explicitly modeling the nonlinear mapping between genotype and phenotype. We defined an altered-specificity PDZ domain variant as one that has a corrected FDR below 0.05 for any substrate. As many of these ligands represent functionally different classes of PDZ ligands, we categorize this dataset as profiling novel substrates or ligands (Class 1).

*Enzymes: TEM-1, AmiE, PafA*

β-lactamases are a class of bacterial serine hydrolases that provide resistance of β-lactam antibiotics such as penicillins and cephalosporins by hydrolyzing the β-lactam ring. TEM-1 is a Class A β-lactamase and the major cause of ampicillin resistance in *E. coli* and other gram-negative bacteria. Because of its easy-to-test phenotype (growth in antibiotic) and evolutionary relevance, it has also long served as a model for studies of enzyme evolution. The most useful such study for our purposes is likely Stiffler et al[48], which both tests complete dose-responses of ampicillin and compares ampicillin resistance to resistance to a third-generation cephalosporin antibiotic, cefotaxime, which TEM-1 natively has low activity against. However, some evolved extended-spectrum β-lactamases do provide resistance against cefotaxime and similar drugs[49], suggesting that this activity is likely represented within the provided MSA, although presumably by a very small fraction of sequences. As cefotaxime is not broken down by TEM-1 and represents a different class of β-lactam antibiotic, we classify this dataset as a profiling new substrates or ligands (Class 1). We based our cutoff for TEM-1 specificity switching on the original Stiffler et al[48] paper: namely with fitness scores against cefotaxime of greater than 0.2.

Aliphatic amidases in prokaryotes are responsible for hydrolyzing various amides to produce ammonia necessary for cell growth. Many of these amidases are quite promiscuous for most aliphatic amides[50]. Wrenbeck and colleagues used one such amidase (AmiE) from *Pseudomonas aeruginosa* which preferentially hydrolyzes very short-chain amides.[50] They tested all single variants of AmiE against three substrates: propionamide and acetamide, both of which AmiE natively hydrolyzes efficiently, and isobutyramide, which is a poor but not fully excluded AmiE substrate. They used a growth-based selection with the amide in question as the sole nitrogen source. As isobutyramide is a (poor) substrate of wildtype AmiE and similar molecules are substrates of closely related amidases, we classify the AmiE dataset as shifting the specificity profile (Class 2). We followed Wrenbeck and colleagues[51] in defining altered-specificity variants as those with fitness scores greater than 0.15 (up to a max of 0.85) for one substrate and less than 0 for the other two substrates.

PafA is a model alkaline phosphatase from the Flavobacterium *Elizabethkingia menangoseptica* known for its extraordinary rate enhancement. Markin et al[52] performed enzyme kinetics measurements on a microfluidic device to profile $k_{cat}$ and $K_m$ values for 1-3 different amino acids at each position of PafA against two phosphate monoesters: methylphosphate (MeP) and 2-(2-oxo-7-phosphonooxy)-2H-chromen-4-yl acetic acid (cMUP; Supplemental Figure 1A-B). Both molecules are substrates of the native enzyme, although the authors propose that they may have different rate-limiting steps: catalysis for cMUP or binding for MeP. Thus, mutations altering the balance between these substrates may in fact reflect differential effects on substrate binding or catalysis. We nonetheless consider this a shift in specificity for existing substrates (Class 2), providing a possible microscopic mechanism for an allosteric specificity mechanism in this example. We determined altered-specificity variants of PafA by fitting a linear trend to the $\log(k_{cat}/K_m)$, summarized in Supplemental Figure 1C-D, with technical details in the methods. Variants with "global catalytic activity" ($g_x$) values below 3.3 $\log(M^{-1}\ s^{-1})$ were deemed inactive, and active mutants with specificity activity ($s$) values above 0.55 $\log(M^{-1}\ s^{-1})$ were deemed to have altered specificity (Supplemental Figure 1C-D), defined based on breaks in the respective distributions.

*Transcription factors: Pho4 and RamR*

Pho4 is a basic helix-loop-helix (bHLH) transcription factor in *S. cerevisiae*, where it regulates phosphate metabolism.[53] Aditham and colleagues used medium-throughput STAMMPing (simultaneous transcription factory affinity measurements via microfluidic protein arrays) to profile the binding affinities of 210 Pho4 mutants against nine variations on Pho4's cognate DNA sequence, 5'-CACGTG-3', and its two flanking nucleotides (C and A).[54] Some DNA sequences have relatively poor binding affinities and represent deviations from Pho4's defined specificity role, and thus we classify the Pho4 dataset in Class 1. We fit altered-specificity variants with linear regression, as for PafA (Supplemental Figure 1E-H).

RamR is a member of the broad TetR family of allosteric ligand-responsive transcription factors in bacteria.[55] This family respond to a diverse array of ligands, many of them unknown.[56] d'Oelsnitz and colleagues tested over 130,000 variants to RamR against three different enantiomeric ligands: 1-DHIQ, 1S-THIQ, and 1R-THIQ.[57] By tying transcriptional depression by

RamR to cell growth in the presence of zeocin, they profiled multiple concentrations of each ligand for each variant, and from these estimated dose-response curves using a Gaussian process model. We used two different concentration levels, 32 μM and 500 μM ligand for our classifications (Supplemental Figure 2). Global deleteriousness was determined based on the fold-induction of transcriptional response at 500 μM ligand, to identify globally inactive variants. We assume that classification as active variant requires the protein to be folded, be able to bind DNA and a ligand (though with almost any affinity), and have an intact allosteric response. We then used the response values at the more stringent condition of 32 μM ligand, selecting only for high affinity, to define altered-specificity variants (Supplemental Figure 2B). Because the wildtype protein only responds to 1S-THIQ and not 1R-THIQ or 1-DHIQ at this concentration, we classify altered-specificity variants as those with a log(relative response) value greater than 0.4. We used this relatively low threshold rather than a more stringent one as no single variants in the library met considerably more stringent thresholds. Based on RamR's strong selection for 1S-THIQ over 1R-THIQ—and the identification of variants with inverted selectivity—we classify the RamR dataset in Class 1.

*Transporters: NorA and DraNramp*

Bacterial multidrug efflux pumps are common drivers of antibiotic resistance,[58] and as such their specificity for different compounds has been a topic of great interest. NorA is a key efflux pump in the core genome of *Staphylococcus aureus* present in all MRSA strains.[59] It is also a member of the massive major facilitator superfamily, which contains transporters with extremely diverse substrate specificities and biochemical properties.[60] Miller and colleagues profiled the extent to which each single mutant of NorA rescued growth in the presence of one of seven small molecule drugs, each of which are natively exported by NorA.[61] We classify the NorA mutational scan in Class 2, as all of the substrates are native to the transporter. Each drug was tested between its minimal inhibitory concentration (MIC) and its $IC_{50}$, meaning that different drugs were presented to the NorA protein for efflux at very different concentrations. We follow Miller and colleagues[61] in using a "specificity index" (the variance of fitness scores between substrates for each position) to define mutations that alter specificity, setting an arbitrary threshold of 2.

Nramps represent a different family of transporters from another one of the largest transporter superfamilies, the APC superfamily sharing a LeuT-fold.[62] These proteins are striking for their remarkable specificity for the import of rare transition metals over common alkaline earth metals such as $Mg^{2+}$ and $Ca^{2+}$,[63] although distant relatives can transport other metals such as $Mg^{2+}$[64] and $Al^{3+}$.[65] We profiled a set of single to octuple mutations of a model bacterial Nramp homolog from the extremophilic bacterium *Deinococcus radiodurans* (*Dra*Nramp) for their ability to import a representative substrate ion, $Mn^{2+}$, and non-substrate ion, $Mg^{2+}$, identifying mutations and combinations of mutations that permit $Mg^{2+}$ import.[66] As with NorA, $Mn^{2+}$ and $Mg^{2+}$ were tested at very different concentrations (100 μM $Mn^{2+}$ and 1 mM $Mg^{2+}$) and assayed differently, although the complete exclusion of $Mg^{2+}$ by wildtype DraNramp means that a variant that is capable of importing any $Mg^{2+}$ at all represents a major specificity shift. We classify the DraNramp study as profiling new substrate or ligands (Class 1).

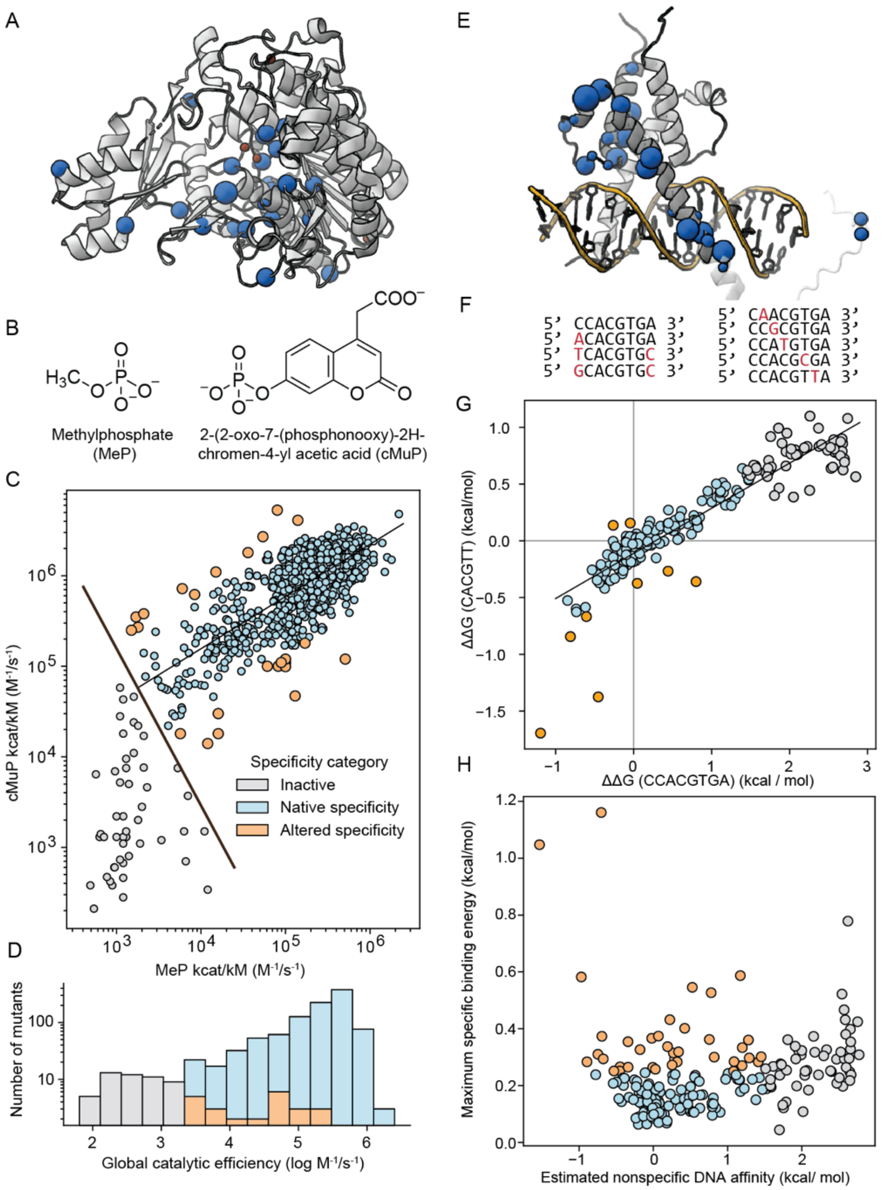


**Supplemental Figure 1. Classification of altered-specificity variants from microfluidics data with linear regression.** (A) Structure of PafA239 (PDB: 5TJ3) with resulting specificity-altering mutations colored, reproduced from Figure 1. (B) Chemical structures of the two PafA substrates tested in Markin et al[52]: methylphosphate (MeP) and 2-(2-oxo-7-phosphonooxy)-2H-chromen-4-yl acetic acid (cMUP). The phosphate group of cMUP is represented as chemically neutral to match the schema in the original paper.[52] (C) Scatterplot of catalytic efficiencies of all tested single variants to PafA for both MeP and cMUP, plotted similarly to Figure 3C of Markin et al.[52] The regression line to active variants of PafA is shown in black, with the perpendicular cutoff line for calling “inactive” variants highlighted in blue. Based on this, inactive variants are shown in gray, native-specificity variants in light blue, and altered-specificity variants in orange. (D) Histogram of the base-10 logarithm of the fit “global” catalytic efficiency of each variant, defined as the relative location of each variant along the regression line (see Methods). Categories are highlighted as in the scatterplot in (C). (E) Structure of Pho4 bound to DNA (PDB: 1A0A)[67] and specificity-altering mutation locations reproduced from Figure 1. (F) The nine DNA sequences tested in

Aditham et al[54]: the cognate sequence 5'-CCACGTGA-3', three variations on the flanking nucleotides, and five mutations to different positions within the core sequence motif. (G) Scatterplot of measured ΔΔG values for the cognate DNA sequence against a representative alternate sequence. As in (C), a regression line and mutation categorizations are displayed, but the global active-inactive threshold is not displayed as this depends on all nine ligands. (H) The estimated nonspecific DNA affinity (equivalent to the global catalytic efficiency in D but defined over eight comparisons) for each variant against the maximum specific binding energy. Note that while the specific binding energy is nondirectional, positive affinity values are worse (higher energy). Mutations are classified using the same color scheme as in (C) and (D).

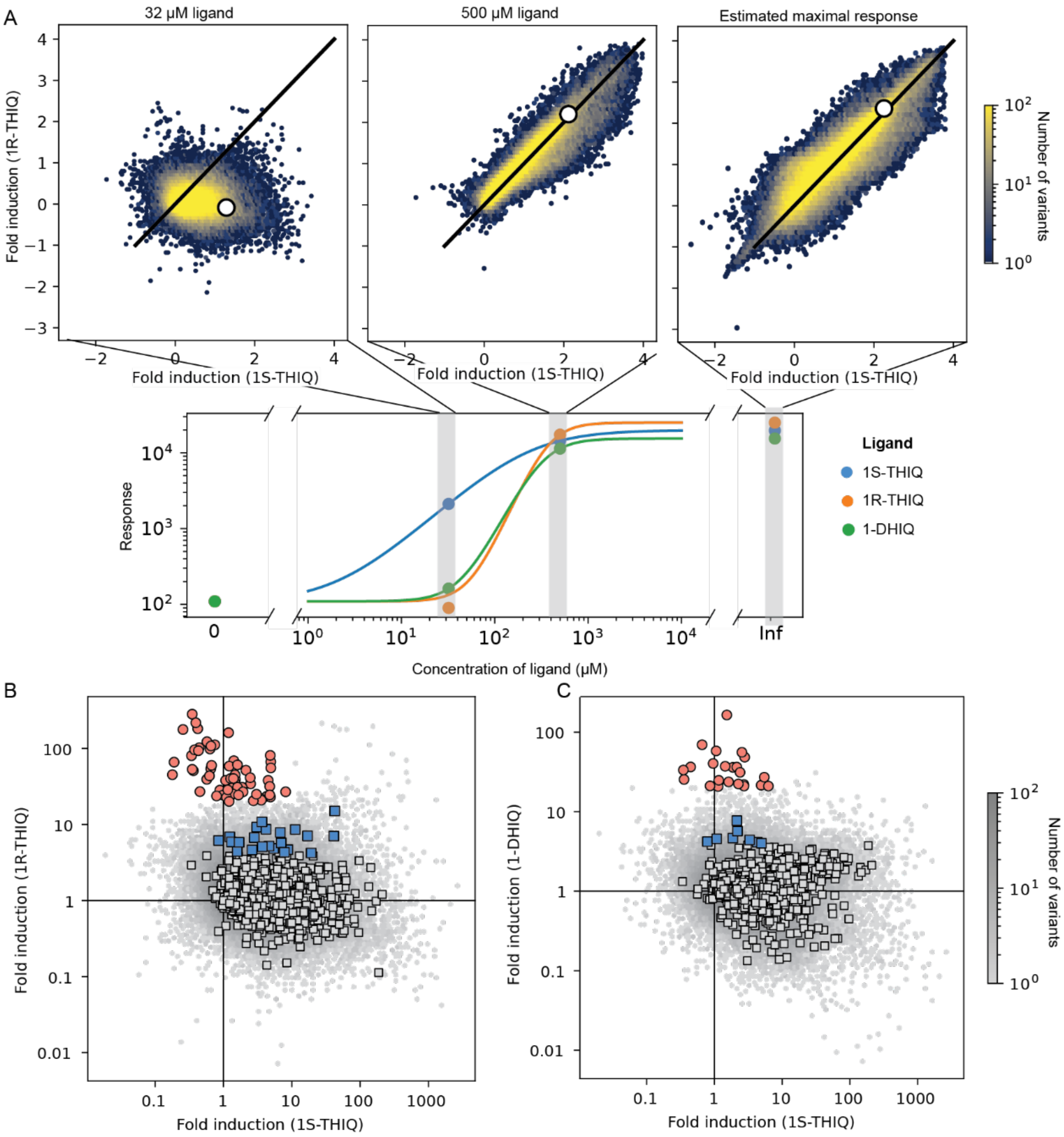


**Supplemental Figure 2. Classification of altered-specificity variants from RamR dose-response curves.** (A) Density plots of fold induction of RamR variants for the enantiomeric ligands 1S- (x-axis) and 1R-THIQ (y-axis) at 32 and 500 µM, as well as the estimated maximal response from a Gaussian process model.[57] Lines of equal response are shown in black and wildtype RamR is highlighted in white. The relative positions of the variants tested on schematized dose-response curves for wildtype RamR are shown below, with points representing actual screen values. At 32 µM, only 1S-THIQ elicits a response, while both ligands elicit similar responses at 500 µM. (B) Distributions of 1S-THIQ and 1R-THIQ fold-induction values at 32 µM, with single mutants highlighted in boxes; those passing the specificity threshold are colored blue. The threshold in question is relatively lenient, representing only a 4-fold induction for the alternate ligand. A more stringent threshold only achieved by multi-mutants is highlighted in red, showing mutants that switch the enantioselectivity of RamR. (C) The same plot for 1S-THIQ against 1-DHIQ with equivalent thresholds.

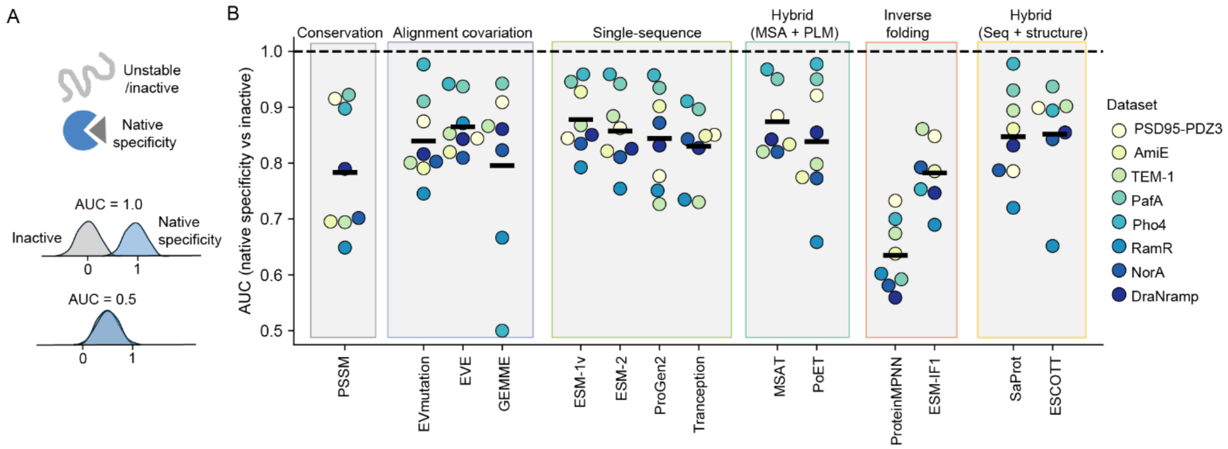


**Supplemental Figure 3. Model performance at separating native-specificity and inactive variants.** The ability of each model to separate variants with native specificity from those deemed globally inactive in each dataset, quantified as the area under the ROC curve (AUC). Models are separated by class, which is labeled above, and individual points are colored by dataset. Horizontal black lines represent mean AUC across datasets for each model.

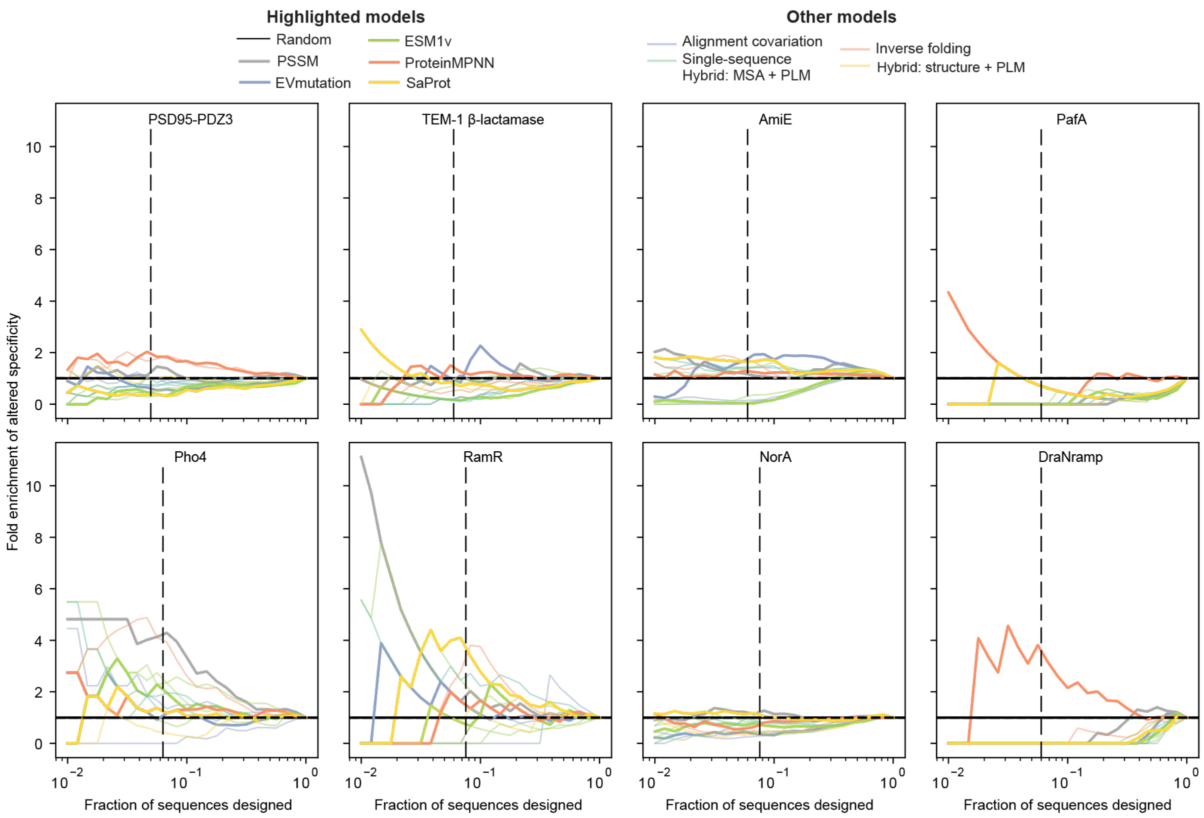


**Supplemental Figure 4. Design enrichment curves for single mutants to all proteins.** Fold-enrichment of altered-specificity variants when varying percentages of top-ranked simulated "designs" among single mutants in each dataset.
